# Garments that measure EEG: Evaluation of an EEG sensor layer fully implemented with smart textiles

**DOI:** 10.1101/2022.12.31.522227

**Authors:** Eduardo López-Larraz, Carlos Escolano, Almudena Robledo, Leyre Morlas, Alexandra Alda, Javier Minguez

## Abstract

This paper presents the first garment capable of measuring EEG activity with accuracy comparable to state-of-the art dry EEG systems. The main innovation is an EEG sensor layer (i.e., the electrodes, the signal transmission, and the cap support) fully implemented as a garment, using threads, fabrics and smart textiles, without relying on any metal or plastic materials. The garment is interfaced via a connector to a mobile EEG amplifier to complete the measurement system. The new EEG system (Garment-EEG) has been characterized with respect to a state-of-the-art Ag/AgCl dry-EEG system (Dry-EEG) over the forehead area of healthy participants in terms of: (1) skin-electrode impedance; (2) electrophysiological measurements (spontaneous and evoked EEG activity); (3) artifacts; and (4) user ergonomics and comfort. The results show that the Garment-EEG system provides comparable recordings to Dry-EEG, but it is more prone to getting affected by artifacts in adverse recording conditions due to poorer contact impedances. Ergonomics and comfort favor the textile-based sensor layer with respect to its metal-based counterpart. User acceptance is the main obstacle for EEG systems to democratize neurotechnology and non-invasive brain-computer interfaces. EEG sensor layers encapsulated in wearables have the potential to enable neurotechnology that is naturally accepted by people in their daily lives. Furthermore, by supporting the EEG implementation in the textile industry it is manufactured with lower cost and much less pollution compared to the metal and plastic industries.

## Introduction

The electroencephalogram (EEG) is a non-invasive technique for measuring the electrical activity produced by the brain (Biasiucci et al., 2019). The firing of neurons (i.e., action potentials) generates electrical currents, and the sum of activations of large groups of neurons spiking synchronously produces voltage differences that can be measured with electrodes placed on the scalp (Niedermeyer and Lopes da Silva, 2005; Nunez and Srinivasan, 2006). The measurement of these neural populations has facilitated the study of human nature, from perception and cognition to action. The EEG is widely used in research and clinical practice. Some of its most relevant applications are for diagnosis of conditions such as epilepsy, ADHD or sleep disorders, among others, and as a brain mapping tool for neuroscience, psychology and electrophysiology research (Niedermeyer and Lopes da Silva, 2005).

One trend of innovation in the EEG field in the last 2 decades addresses the evolution of EEG technology to enable its use for real-world research and real-life applications. This has led to the development of wireless systems, their miniaturization, low-density sensor layouts, the use of new conductive materials to produce gel-less sensors, improved usability for non-experts, and cost reduction, among many other innovations (Casson et al., 2010; Millán et al., 2010; Casson, 2019). All these efforts are directed towards the improvement of user acceptance, while trying to preserve signal quality. Although there has been very relevant progress in this direction, reliable EEG technologies that are affordable and widely accepted by the general population are still missing.

Mobile EEG systems are composed by three elements: 1) the EEG sensor layer (i.e., EEG electrodes, signal transmission and head support); 2) the EEG amplifier (analogic, digital and communication electronics); and 3) the interface between them (connector), see Figure 1. The sensor layer is the component that contributes most to the usability/ergonomics of the whole system, and thus, is key for its acceptability in out-of-the-lab environments (Biasiucci et al., 2019; Casson, 2019). This layer, and specifically the EEG electrodes, traditionally rely on wet electrodes (Figure 1, first row), which require the application of an electrolytic substance between the sensor and the scalp. Although they are the gold-standard in medical applications, this reduces their usability and acceptance. Dry-EEG electrodes do not require the application of electrolytes (Figure 1, second row), which improves setup, cleaning time, ergonomics and comfort, especially for non-experts (Lopez-Gordo et al., 2014). These dry electrodes operate with higher contact impedances and thus are more prone to artifacts than wet electrodes, which is partially compensated by higher input impedance amplifiers (Shad et al., 2020). There has been a lot of research in new conductive materials and electrode shapes to create novel dry-EEG electrodes: different conductive metals and coatings (Lopez-Gordo et al., 2014), hydrogels (Li et al., 2021; Pei et al., 2022), polymers (Leleux et al., 2014), conductive inks (Ferrari et al., 2020), graphene (Ko et al., 2021), organic transistors (Venkatraman et al., 2018), conductive/smart textiles (Tseghai et al., 2020), among others.

**Figure 1.**
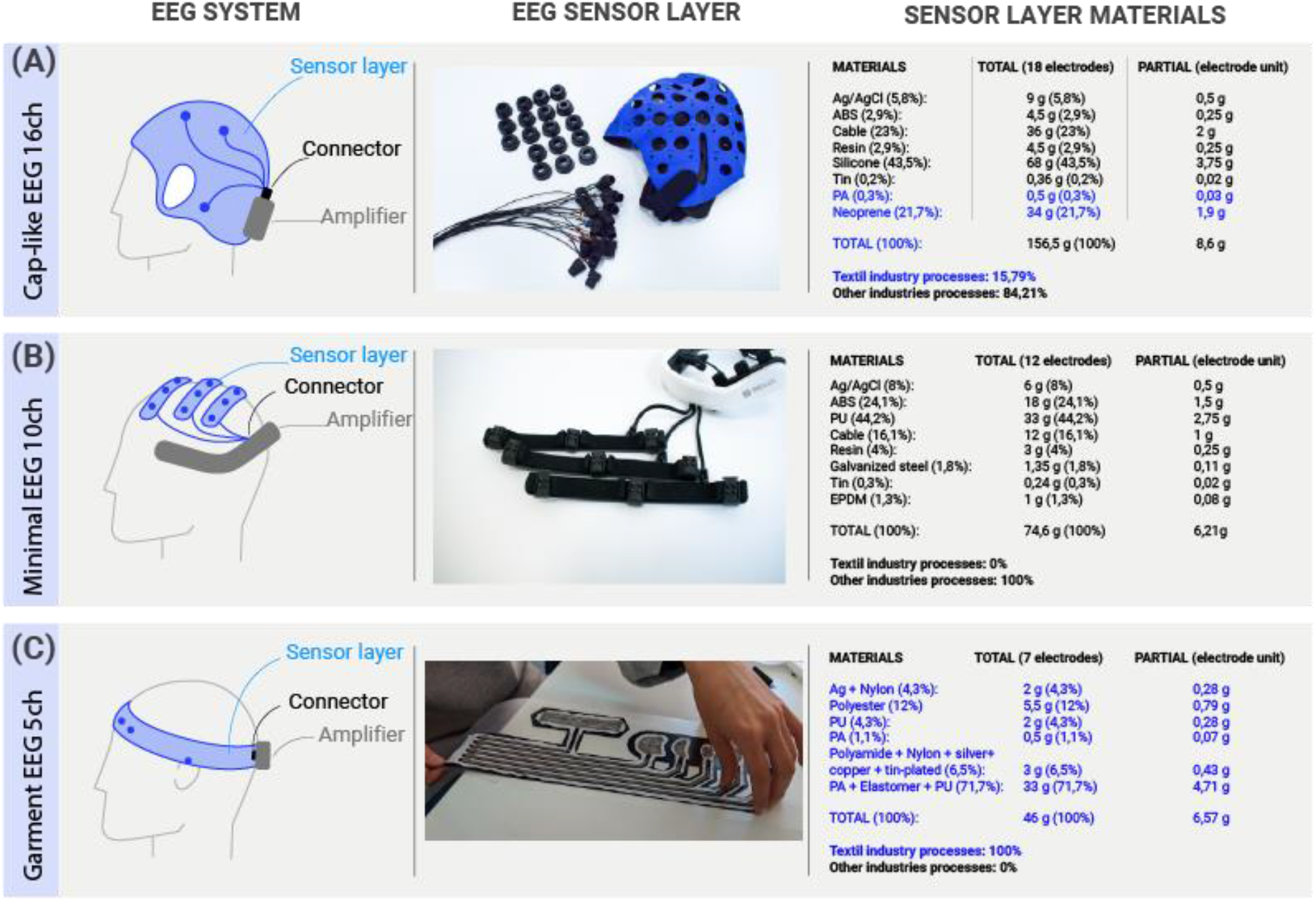
Different technologies for EEG devices with specific focus on the EEG sensor layer. (A) Example of “Shower-cap” research-oriented EEG systems. (B) Example of “Minimalistic” application-oriented EEG systems. (C) Proof-of-concept “Garment” EEG system. In all the EEG devices, we differentiate between the EEG sensor layer, the connector, and the amplifier. In A and B devices, the sensor layer is implemented with plastics, metals, glues and materials from the electronic devices industries, while in C, “Garment” sensor layers are implemented with materials and manufacturing processes from the textile industry only. The blue text in the right panel represents materials and manufacturing processes from the textile industry.

Conductive or smart textiles are especially advantageous for the development of EEG sensors (usually referred to as textile-EEG or textrodes), but also to completely integrate the EEG sensor layer as a garment (for their differentiation we will denote them as “Garment-EEG”). These sensors or garments can be built by combining conductive and isolating fibers, being light, soft and, bendable, which increases the comfort for the user and reduces their fabrication cost. They can also inherit some relevant properties from the textile industry, like being breathable and washable. There are successful examples of biosignal electrodes based on textiles (Márquez et al., 2013) to measure ExG signals within the millivolt range, like electrocardiogram (ECG), electromyogram (EMG) and electrooculogram (EOG) (Catrysse et al., 2003; Acar et al., 2019). However, the EEG is measured in the microvolt range, where obtaining an acceptable signal-to-noise ratio is far more challenging. Note that, while standard wet EEG has impedances below 5 KΩ, textile (and other dry) sensors are two orders of magnitude above (in the range of 100s of KΩ), and thus, have higher levels of noise. To date, only a few research studies have proposed innovative EEG electrodes with different conductive textile materials, such as silver-plated or copper-plated fabrics (Tseghai et al., 2020). A recent review of the literature on textile EEG suggested that the lack of relevant progress in this field is mainly due to 2 factors. On the one hand, the new textile EEG systems are limited to publishable research, with no incremental or further innovative development. On the other hand, these published developments do not generally report quantitative comparisons with standard EEG systems to assess their value (Tseghai et al., 2020). When generalizing from the electrode to the entire sensor layer, all proposed textile EEG systems always rely on non-textile materials (metals or plastics) to complete the sensors, transmissions, or cap support. For instance, by developing textile electrodes that are used to contact standard metal electrodes (Fleury et al., 2017), or by connecting the textile electrodes via metal snaps to standard cables (Löfhede et al., 2012; Arnin et al., 2013).

In this paper, we propose the proof of concept of an EEG sensor layer implemented using only materials and manufacturing processes from the textile industry (Figure 1, third row). In other words, the complete sensor layer is a garment by design and manufacture. The contribution is the unified design of the EEG electrodes, the shielded transmission and the head support using conductive and isolating textile materials. The new EEG system (Garment-EEG) has been quantitatively characterized with respect to a state-of-the-art Ag/AgCl dry EEG system (Dry-EEG) over the forehead area of healthy participants in terms of: 1) skin-electrode impedance; 2) electrophysiological measurements (spontaneous and evoked EEG activity); 3) artifacts; and 4) user ergonomics and comfort.

## Materials and Methods

To characterize the new EEG sensor layer, we designed two different headbands to monitor the EEG from people’s forehead, at positions F7, Fp1, Fp2 and F8, according to the International 10/20 system. One of them was made exclusively with textile materials (Garment-EEG) and the other integrating standard metal electrodes and coaxial cables (Dry-EEG). This Dry-EEG technology been selected as baseline and state-of-the art for the comparison as it has been already benchmarked with respect to a medical-grade gel-EEG system in the context of electrophysiological measurements and brain-computer interfaces (Schwarz et al., 2020). Both headbands included 4 recording electrodes, plus reference and ground electrodes. They had the same type of connector at their back, where the amplifier was connected. The layout simplification covers the frontal brain area to simplify the technology and focus on a brain area with easy access and replicable setup. This allows to focus on the skin-electrode properties and quality of the electrophysiological data, avoiding artifacts derived from contact with hairy areas (placement, difference type of hairs, head morphology, etc.).

### Garment-EEG headband

The novel Garment-EEG headband integrated the sensors and the signal transmission using only textile materials (Figure 2A). The textrodes were implemented using 3-strand silver-coated Nylon conductive yarns with a linear resistance of 114 Ω/m. They were embroidered in 3 directions (0º, - 45º, and 45º) with a distance between yarns of 1mm. Electrodes at Fp1 and Fp2 covered an area of 2.8 cm^2^, while F7 and F8 electrodes covered an area of 7.4 cm^2^, to allow an appropriate electrode-skin contact for different head sizes. The ground electrode was placed on Fpz and covered an area of 1.5 cm^2^. The reference electrode had a long rectangular shape of 18 cm^2^ to facilitate the contact on the upper union between the left ear and the head regardless of the head perimeter.

**Figure 2.**
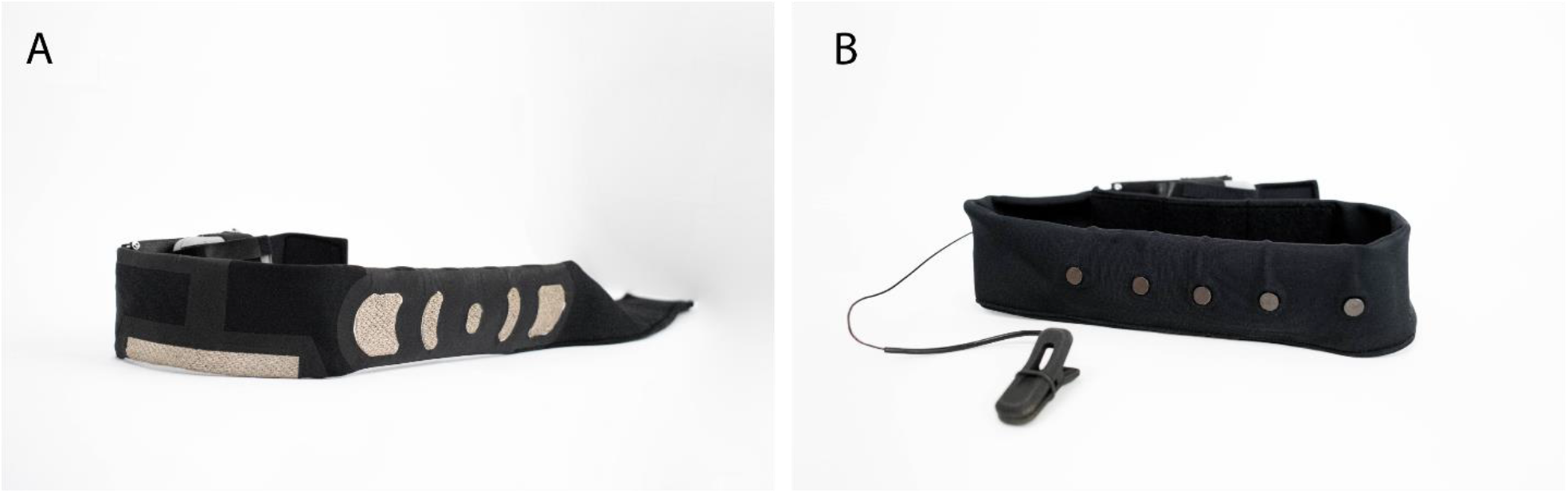
Garment-EEG and Dry-EEG headbands. (A) Detail of Garment-EEG electrodes shape and location. (B) Detail of Dry-EEG electrodes shape and location.

The signal transmission from the sensor to the connector was implemented mimicking a coaxial cable with active shielding, by overlapping different textile layers. First, the transmission wire was embroidered with 8-strand silver-coated Nylon yarns, with a linear resistance of 15 Ω/m. This conductor was insulated using vinyl by heat-sealing at a temperature of 150ºC for 10 seconds. Then, an additional conductive layer was added to shield both sides, using Ultraflex Tape Zell RS conductive fabric, made of PA/Nylon 6.6 coated with silver, copper, and tin, and with surface resistivity of less than 0.02 Ω. The final overlay, to isolate and fix the transmission to the fabric was made with vinyl, sealed at 150°C for 10 seconds.

The connection between the textrode and the transmission core was made by embroidering the former directly onto the latter. The connection between the transmission and the connector was made through a rigid PCB with copper pads sewn with conductive wire made of silver-coated polyamide/polyester, with a linear resistance of less than 530 Ω/m. The support headband was made of polyester fabric with a satin structure and a grammage of 120g/m2.

### Dry-EEG headband

The Dry-EEG system integrated Ag/AgCl electrodes, welded on coaxial cables, onto a similar headband (Figure 2B). The 4 recording electrodes were located at F7, Fp1, Fp2 and F8 positions. The ground electrode was placed on Fpz, and the reference electrode was on the left ear with an earclip. These electrodes had a circular shape with a diameter of 0.8 cm. The cables included a silver platted copper core (0.45 mm of diameter, resistance of 0.158 Ω/m), a PTFE tape insulation, a silver platted copper shielding layer, and an extruded FEP jacket. The connection between the coaxial cables and the connector was made by welding. The support headband was made of polyester fabric with a satin structure and a grammage of 120g/m2 (similar to the textile one).

### EEG connector and amplifier

The same biosignal amplifier was used with the two EEG headbands. It was a high input impedance amplifier (10^4^ GΩ / 5 pF) with a CMRR of 100 dB, to reduce the noise and the imbalance with respect to the skin-electrode impedance (in dry-EEG, impedances can be in the order of 100-200KΩ, see (Lopez-Gordo et al., 2014; Shad et al., 2020)). The system provides active shielding for the sensors and a DRL circuit to reduce noise. This amplifier samples the 4 EEG channels at 256 Hz and uses Bluetooth Low Energy (BLE) to transmit the data to a laptop. To connect the sensor layer and the amplifier, we used a commercial 20-position board-to-board connector with gold-plated phosphor bronze contacts and a liquid crystal polymer cover.

### Experimental validation

We carried out two experimental studies. In the first one, we characterized the impedance of the electrodes and signal transmission of the new Garment-EEG system, while in the second, we quantified common spontaneous (frequency domain) and evoked (time-frequency domain) EEG patterns of activity.

### Participants

Six healthy volunteers participated in the first study (5 females, 1 male, age: 30.2±5.8 years) and ten different healthy volunteers participated in the second study (5 females, 5 males, age: 27.8±3.7 years). The participants were recruited from Bitbrain’s volunteer database and received an economic compensation of 25€. All of them signed an informed consent form and were debriefed about the study purpose and their rights regarding the physiological and demographic data that were collected from them.

### Study 1: Impedance analysis

We quantified the impedances of the sensor layer (including the electrode and the transmission) of both the Garment-EEG and the Dry-EEG systems. As baseline, we performed an additional measurement (Wet-EEG) in which the Dry-EEG headband was used with an electrolytic substance (SAC2 conductive gel, by Spes Medica) after cleaning the skin with abrasive gel (NeurPrep, by Spes Medica). This skin preparation with abrasive gel is a common procedure in clinical and research EEG applications. To simulate real-world conditions in the evaluation of the Garment-EEG and Dry-EEG impedances, the skin of the subjects was only cleansed with make-up remover wipes.

We used a PalmSens4 with a MUX8-R2 multiplexer (PalmSens, Houten, Netherlands) to measure the skin-electrode impedance as a function of logarithmically sampled frequency, ranging from 0.1 Hz to 30 Hz with 10 sample points. We repeated the impedance measurements 3 times for each of the 4 electrodes to obtain more reliable estimations.

### Study 2: EEG activity analysis

In this study, the brain activity of the participants was monitored using, sequentially, the two headbands. They performed the same tasks using both technologies in a cross-over manner, half of them starting with the Garment-EEG and the other half starting with the Dry-EEG (serial setup for EEG comparisons, see (Gargiulo et al., 2010; Kutafina et al., 2020)). The tasks were:

- **Task 1**: 3 minutes of resting state with eyes closed.
- **Task 2**: 3 minutes of resting state with eyes open.
- **Task 3**: 80 cue-guided reaching movements with the right arm (divided in 4 blocks of 20 movements each), which included a resting period of 7±1 s, and a movement period of 5 s.
- **Task 4**: artifact induction, where the following actions were executed for 10 seconds each, to contaminate the EEG signals: *i)* the experimenter moves both hands above the participant’s head; *ii)* tongue movements; *iii)* jaw movements; *iv)* blinking; *v)* lateral and vertical eye movement; *vi)* vertical head movement; *vii)* horizontal head movement; *viii)* up and down shoulder movement; *ix)* up and down movement with both arms.

### EEG analysis

The EEG signals recorded with both technologies were quantified and compared using a frequency analysis (tasks 1, 2 and 4) and a time-frequency analysis (task 3).

### Frequency analysis of EEG during resting state/artifact induction (Tasks 1, 2 and 4)

The eyes-closed and eyes-open resting state (Tasks 1 and 2), and the artifact induction (Task 4) EEG recordings were analyzed in terms of their power spectral density.

#### Preprocessing

First, we removed outlier amplitude values from each EEG recording. For that, we high-pass filtered the signals at 60 Hz, with a 4-th order non-causal Butterworth filter, applied a Hilbert transform, and z-scored the values. Those samples going beyond 5 standard deviations above the mean were marked as outliers and discarded in the original signal, filling the gaps by linear interpolation. This step was not applied to the signals of the artifact induction task, since in that case, the objective was to compare the occurrence of artifacts in both technologies.

#### Processing

We filtered the EEG signals between 0.1 and 100 Hz, using a 4-th order non-causal Butterworth filter. Subsequently, we estimated their power distribution between 0 and 60 Hz using a fast Fourier transform (FFT) with a 1 s hamming window, zero-padded to 1024 points (0.25 Hz resolution) and 30 ms of overlap, and computed the logarithm of the obtained power values.

#### Statistics

For the statistical comparisons, we averaged the power values of the four electrodes. For this *average* electrode, we computed the mean power for each participant and headband in five frequency bands: 0-3 Hz (delta), 4-7 Hz (theta), 8-12 Hz (alpha), 13-30 Hz (beta), and 45-55 Hz (electrical noise). These ranges cover the most common frequency bands in EEG analyses, as well as the influence of electrical contamination, which has been used as a surrogate of electrode impedance (Insausti-Delgado et al., 2021). A paired Wilcoxon signed-rank test was used to test for statistical differences between both headbands in each frequency band.

### Time-frequency analysis of EEG during movement execution (Task 3)

The EEG data recorded during the execution of reaching movements was analyzed in the time-frequency domain to quantify the event-related desynchronization/synchronization (ERD/ERS) of sensorimotor rhythms (Pfurtscheller and Lopes da Silva, 1999).

#### Preprocessing

First, we removed trials that contained unusually high amplitudes. Trials showing amplitude values higher than 250 μV in any of the four electrodes were marked as artifacts and discarded.

#### Processing

We filtered the EEG signals between 0.3 and 30 Hz, using a 4-th order non-causal Butterworth filter. Subsequently, we computed the time-frequency representation of all the trials using Morlet Wavelets in the frequency range 5-30 Hz, with a frequency resolution of 0.25 Hz. The ERD/ERS maps were computed as the percentage of power decrease/increase with respect to a resting state baseline ([-5, -2] s).

#### Statistics

To compare the ERD/ERS obtained with the Garment-EEG and the Dry-EEG headbands, we first averaged the power values of the four electrodes. We pooled together the trials of all the subjects for each headband and compared them in each time-frequency bin of the *average* electrode using a Mann–Whitney U test.

### Comfort and perception assessment

After completion of the EEG recordings with each sensor layer, participants filled a comfort and perception questionnaire, in which they were asked about the following four points:

- Global comfort of the system, including the electrodes, the headband and the amplifier, on a scale from 1 (“Not comfortable at all”) to 7 (“Very comfortable”).
- Comfort of the EEG electrodes and their contact with the forehead skin, on a scale from 1 (“Not comfortable at all”) to 7 (“Very comfortable”).
- Weight perception of the system, using the following scale: 1 (“I do not feel the weight”), 2 “Noticeable but not bothersome”, 3 “Noticeable but little bothersome”, and 4 “Very noticeable and bothersome”.
- Perception of stability of the system, using the following scale: 1 “I feel it stable”, 2 “I feel it a bit unstable”, 3 “I feel it unstable”.

## Results

### Impedance analysis

Figure 3 shows the impedance values as a function of the frequency for the Garment-EEG, Dry-EEG and Wet-EEG (Dry-EEG using electrolytic gel). The average impedances at 1 and 10 Hz for the Garment-EEG were 368 and 171KΩ, for the Dry-EEG were 110 and 78KΩ, and for the Wet-EEG were 3.2 and 3.05KΩ (below the gold-standard threshold of 5KΩ). Notice that Garment-EEG has systematically higher impedances than Dry-EEG (around two-three times more), and both of them are two orders of magnitude above Wet-EEG. These results are in line with state-of-the-art values reported for wet and dry EEG systems (Lopez-Gordo et al., 2014; Shad et al., 2020).

**Figure 3.**
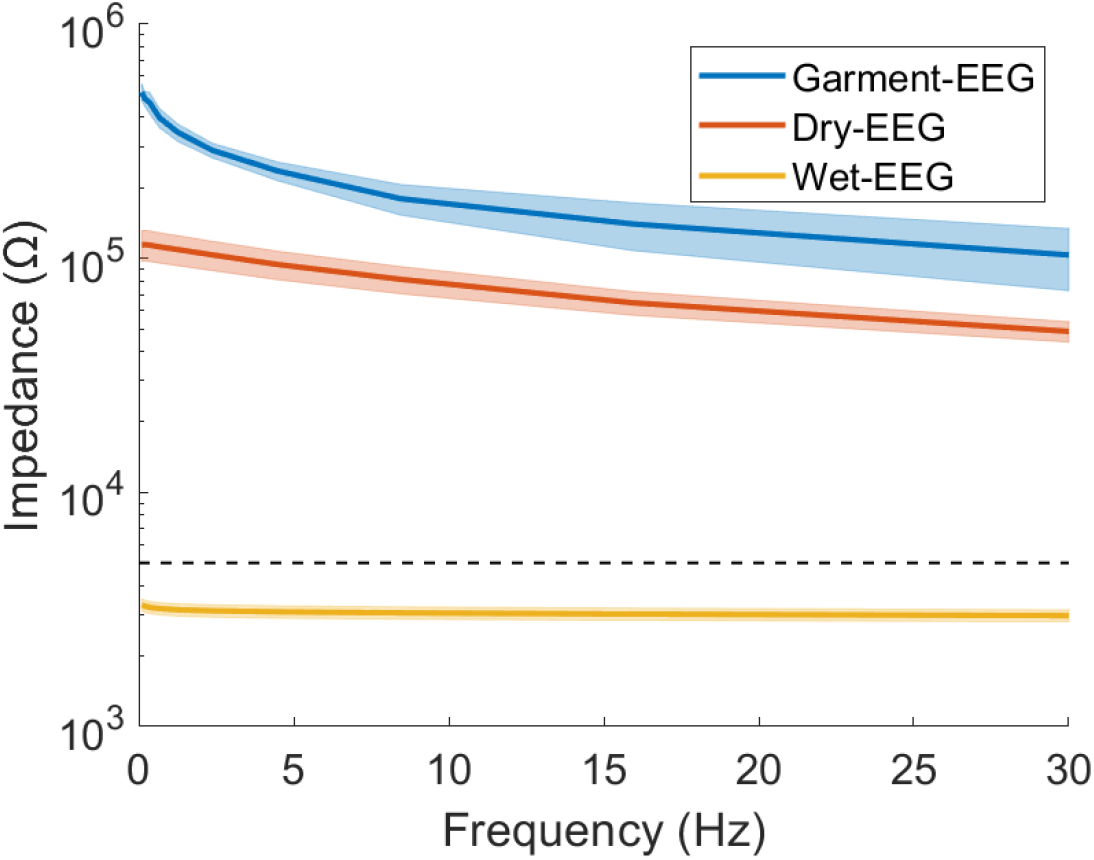
Impedance results for Garment-EEG, Dry-EEG and Wet-EEG. Impedance is displayed as a function of the frequency. The values for all the subjects, electrodes and repeated measurements for each electrode are averaged for each technology. The shades represent the standard error of the mean. Notice that the y-axis is displayed in a logarithmic scale. The dashed horizontal line displays the value of 5KΩ, as standard threshold for good impedance in wet EEG systems (Nuwer et al., 1998).

### Frequency analysis of EEG during resting state/artifact induction

Figure 4 displays the power spectral density analysis of the EEG signals recorded with both headbands during eyes-closed resting state (Figure 4A), eyes-open resting state (Figure 4B), and during the generation of artifacts (Figure 4C). Overall, the signals measured during resting state with the Garment-EEG and the Dry-EEG headbands displayed equivalent power spectral density values at the frequencies usually analyzed in EEG studies (i.e., below 30 Hz). During eyes-closed resting state, a clear alpha peak at ∼10 Hz can be observed in frontopolar (Fp1 and F2) and frontal (F7 and F8) locations using both technologies (Figure 4A). This alpha activity is not so evident during eyes-open (Figure 4B). Noticeably, the Garment-EEG headband resulted to be more prone to getting affected by artifacts. This can be seen as higher power at 50 Hz in all the channels (Figure 4A-B), as well as higher broadband power during the artifact induction task (Figure 4C). For both resting state measurements, the EEG power in delta, theta, alpha and beta frequencies was not significantly different between the Garment-EEG and the Dry-EEG (Figure 4D-E; p>0.05 in all cases). In contrast, the power in these frequencies was significantly higher for the Garment-EEG headband than for the Dry-EEG headband during the induction of artifacts EEG (Figure 4F; p<0.01 in all cases). For the 45-55 Hz frequency band, the power was significantly higher with the Garment-EEG headband both during resting state and during artifact induction (Figure 4D-F; p<0.01 in all cases).

**Figure 4.**
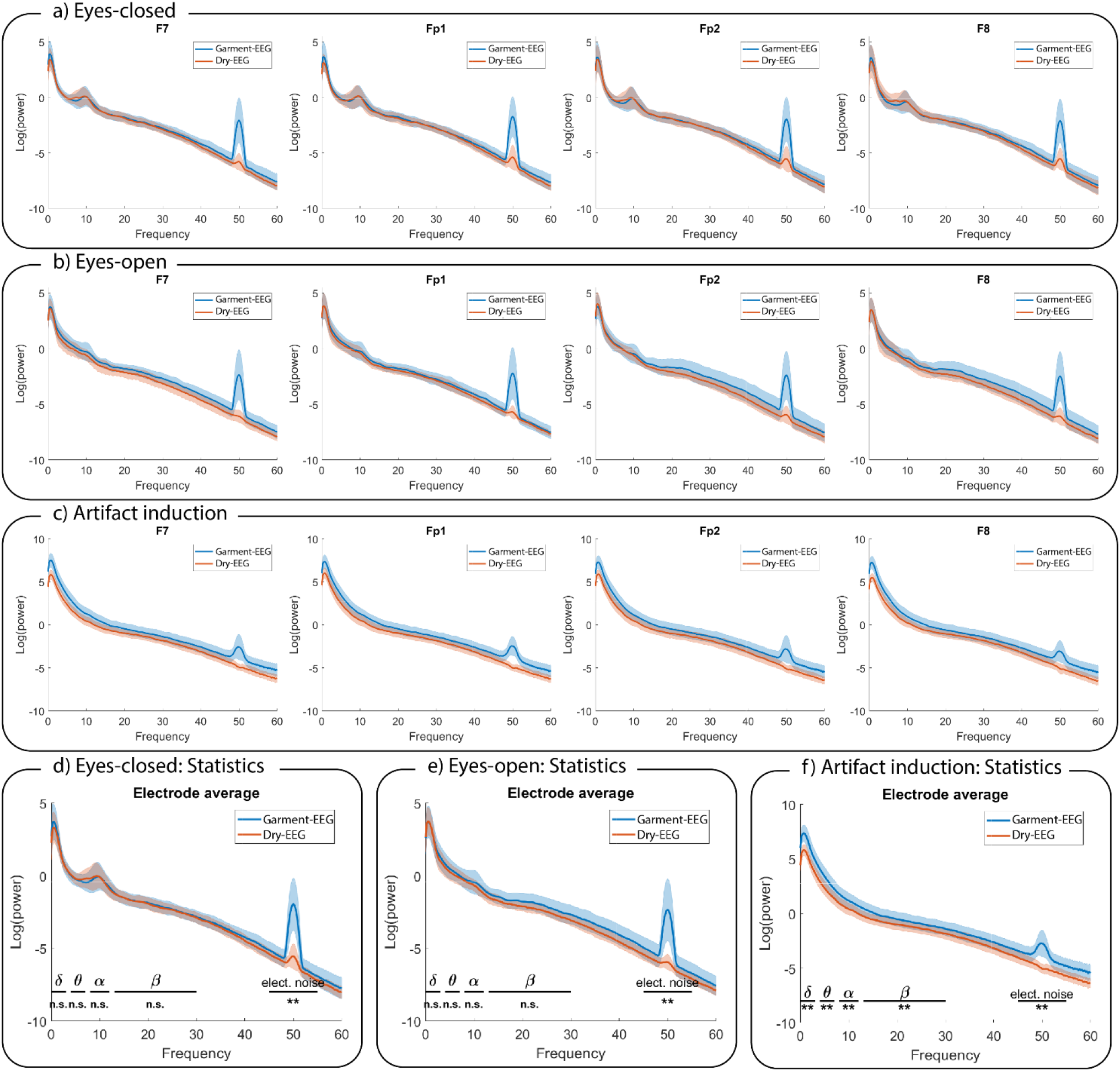
Frequency analysis. Power spectral density comparison between the EEG activity measured with the Garment-EEG and Dry-EEG during (A) eyes-closed, (B) eyes-open, and (C) artifact induction. In panels A-C, each subplot displays one of the four forehead electrodes (F7, Fp1, Fp2, and F8). Statistical comparisons between both headbands on five frequency bands during (D) eyes-closed, (E) eyes-open, and (F) artifact induction. n.s. = non-significant; ** p<0.01.

### Time-frequency analysis of EEG during movement execution

Figure 5 shows the ERD/ERS maps of the four electrodes with the Garment-EEG headband (Figure 5A) and with the Dry-EEG headband (Figure 5B). Given the distant location of the four electrodes with respect to the motor cortex, which is where this motor-related activity originates, the ERD of sensorimotor rhythms is of low magnitude with both technologies. When directly comparing the ERD magnitude of the *average* electrode of both systems, the Dry-EEG headband produces a stronger ERD (Figure 5C, left and center panels). The time-frequency bins that showed a significant difference between both headbands (p<0.01) were used to create a mask that was subsequently applied to the difference between the ERD/S map of the Dry-EEG headband and the ERD/S map of the Garment-EEG headband (Figure 5C, right panel). This analysis revealed that the Dry-EEG headband provided an ERD significantly stronger between 5 and 15 Hz during the execution of the reaching movements.

**Figure 5.**
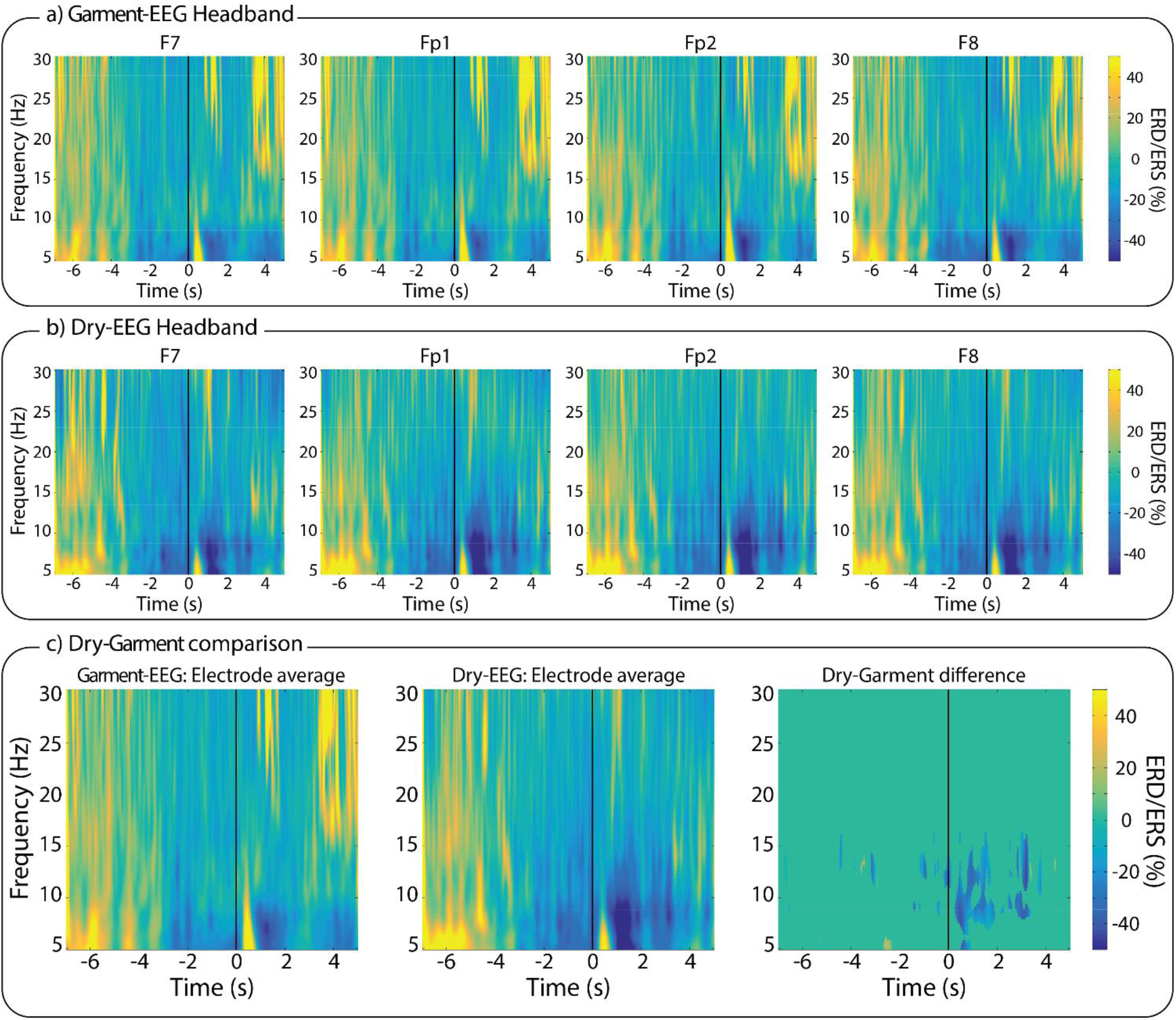
Time-frequency analysis. Event-related (de)synchronization (ERD/ERS) maps of the EEG activity measured during the reaching movements with (A) the Garment-EEG headband, and (B) the Dry-EEG headband. Each subplot in panels A-B displays one of the four forehead electrodes (F7, Fp1, Fp2, and F8). (C) Comparison between the two technologies; (left) average of all the Garment-EEG electrodes, (center) average of all the Dry-EEG electrodes, (right) pairwise statistical comparison between both headbands.

### Comfort metrics

The time the participants spent with each headband ranged between 42 and 48 minutes, except for one case (Participant 1, session with Garment-EEG headband) where, due to technical problems, the session lasted 66 minutes (Figure 6A). The degree of comfort reported by the participants, both for the whole system and for the sensors part, was always high for the Garment-EEG headband and medium-to-high for the Dry-EEG headband (Figure 6B-C). The perception of the weight was distributed between “I do not feel the weight” and “Noticeable but little bothersome”, and the participants never reported any of the headbands to be bothersome (Figure 6D). The perception of stability was also positive, with most of the responses being “I feel it stable”, and none reporting the systems to be unstable (Figure 6E).

**Figure 6.**
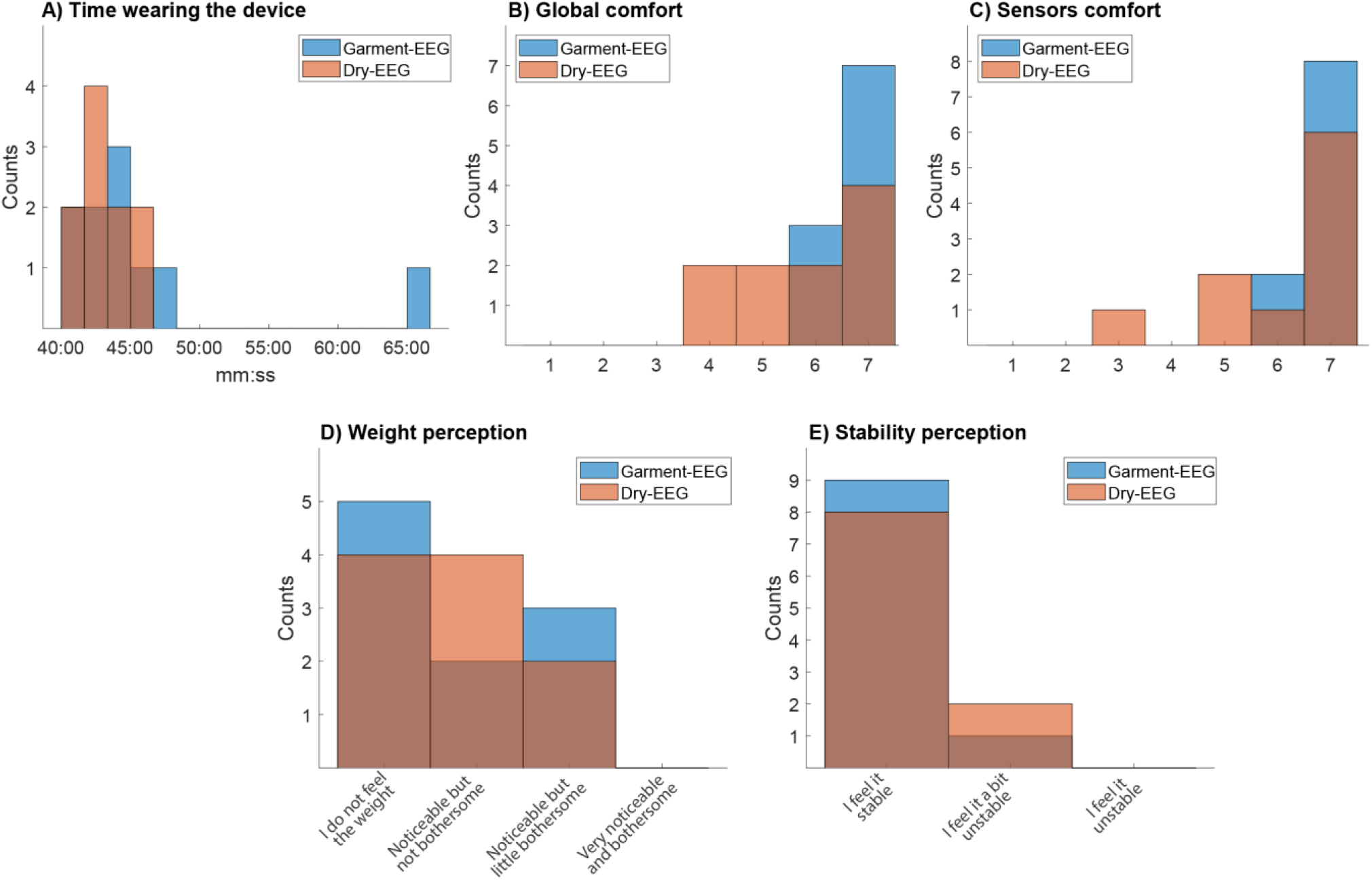
Comfort metrics. Histograms representing (A) the total time the subjects wore the headband, and subjective metrics, such as: (B) global comfort of the headband, (C) comfort of the sensors, (D) perception of the systems’ weight and (E) perception of the system’s stability

## Discussion

Accessible electroencephalographic (EEG) systems are the first step towards democratizing neurotechnology. Wearable market indicators predict a massive adoption of wearable neurotechnology in the next one or two decades, similar to what was experienced a several years ago with other wearables (Johnson and Picard, 2020). Recent hardware innovations in dry EEG technologies have widened the portfolio of tools available for monitoring the brain. However, despite the progress over the last decade, we cannot yet consider that dry electrodes have completely revolutionized the way EEG is used.

User acceptance in terms of ergonomics, usability and price are the main obstacles for EEG systems to establish neurotechnology and non-invasive brain-computer interfaces as a common technology. In principle, EEG sensor layers built with textiles and encapsulated in garments have the potential to enable neurotechnology that is naturally accepted by people in their daily lives. In addition to this, by supporting the EEG implementation in the textile industry, it is manufactured at a lower cost and much less pollution compared to the metal and plastic industries. In this paper, we propose and quantitatively characterize a proof of concept of a garment to measure EEG; a system that uses, exclusively, textile materials for its sensor layer. The evolution and optimization of this technology might open the door to its application out of scientific laboratories or clinics, either for real-world research, for home-based wellness applications, or for patients’ diagnostics, treatment and follow-up.

### Signal quality and artifacts

Overall, the proof-of-concept Garment-EEG proposed in this study provided EEG recordings that were comparable to those of state-of-the-art, metal-based dry EEG electrodes. In terms of the resting state recordings with eyes open and eyes closed, the signals in the frequency domain had equivalent power distributions for both technologies, and there were no significant differences in power between the Garment-EEG and the Dry-EEG at frequencies below 30 Hz (i.e., the ranges usually analyzed in EEG applications). This includes the alpha waves visible in the eyes closed condition and its blocking when eyes are open (Klimesch, 1999).

The motor-related evoked activity produced by arm movements, measured as time-frequency ERD/ERS maps, was visible with both systems over the measured frontal areas, but with a considerably lower magnitude than what is normally measured at the motor cortex during this type of tasks (López-Larraz et al., 2014). Dry-EEG provided slightly stronger ERD maps than the Garment-EEG. Although these results might not be conclusive, there are two possible causes: 1) it could be that artifacts produced during the execution of upper limb movements (Bibián et al., 2022) affect the recording with textile electrodes more than with dry electrodes; and 2) the reference of the Garment-EEG is in the upper part of the ear, being closer to the sensory-motor area than in the Dry-EEG, which could lead to an attenuation of the ERD map.

Garment-EEG was more prone to getting affected by artifacts, which could be seen as significantly higher 50 Hz contamination and significantly higher broadband power during the artifact induction task. Artifacts are one of the most limiting problems in EEG research (Urigüen and Garcia-Zapirain, 2015; López-Larraz et al., 2018), and therefore, the higher likelihood of Garment-EEG of getting contaminated should not be disregarded. However, having more frequent artifacts does not invalidate the relevance of the technology. Artifact removal is a standard procedure (and even a necessary step) in most clinical and research EEG applications (Urigüen and Garcia-Zapirain, 2015), and recent research is currently targeting specific techniques for low density EEG (Lopez et al., 2022). Therefore, while there should be lines of work trying to minimize the appearance and impact of artifacts in this new technology, there can also be progress in the field under the assumption that this type of technology can overcome the limitation of lower data quality with the effortless obtention of massive amounts of data and the exploitation of advanced machine learning techniques (Craik et al., 2019). Note also that this affects recordings in the presence of artifacts, but the fact that signals were comparable in less artifacted conditions (i.e., resting state recordings) is a motivating result that supports further research to continue improving this technology.

One of the factors that could contribute most to the difference in how artifacts affected the Garment-EEG compared with the Dry-EEG is their contact impedance. Impedances of Dry-EEG electrodes have been reported to be in the range of 100-200 KΩ over non-hairy areas (Lopez-Gordo et al., 2014; Shad et al., 2020). This is the reason why high input impedance amplifiers, like the one used in this study, are required (preferably in the order of GΩ). As could be seen, the impedances of the Garment-EEG and the Dry-EEG systems were consistent with those values, although they were around two-three times higher with the textile-based system. Conductive textiles cannot yet offer resistance values at the level of metals, and the fact that both the electrode and the transmission line of the Garment-EEG were made with textiles justifies these higher impedance values. Strategies to optimize the electrode-skin contact (Soroudi et al., 2019; Song et al., 2020) or to maximize the efficacy of the transmission lines (Leśnikowski, 2011) might be a reasonable starting point to continue optimizing this technology.

### Applications of garments to measure EEG

The use of smart textiles to manufacture EEG devices is a growing trend with the potential of large end-user acceptance and lower costs. One of the main critics pointed out by a recent review in Textile-EEG is that most of the systems developed in the last few years are not further developed and remain only as published papers, without a clear progress towards usable and marketable devices (Tseghai et al., 2020). The authors also evidenced a lack of publications with quantitative analysis and from industrial research, which slows the progress in the field (Tseghai et al., 2020). We believe that public-private investment is key to foster the development of radically new technologies such as the one proposed in this study. Once this type of technology reaches an appropriate level of maturity, there is a vast range of applications that could easily benefit from having EEG devices integrated as garments. These could be divided into three contexts: medical, research and wellness/leisure.

While the use of neurotechnology to allow brain control of devices out of the laboratory (e.g., videogames, domotics or even rehabilitation) might still require several years of research and innovation to become a reality, ambulatory brain monitoring is a promising near-term medical application, for instance, for patient follow-up, seizure detection or sleep assessment (Cannard et al., 2020; Tseghai et al., 2020). In sleep studies, it could be used as a relatively cheap triage tool for patients that require a full polysomnographic study (Rundo and Downey, 2019). Epileptic patients can also require long term monitoring of the brain during daily live activities, and it could be easily done with a wearable textile EEG device (Hasan and Tatum, 2021). For the monitorization of newborns, or for patients in need of intensive care, brain monitoring with soft systems like textiles would also constitute a clear advantage (Löfhede et al., 2012).

In the mid-to-long term, this new technology in combination with modern artificial intelligence techniques, could be used to discover and track biomarkers related with the progress or treatment of neurological disorders, such as cognitive decline (Huggins et al., 2021), Parkinson’s disease (Scholten et al., 2020), stroke (Ray et al., 2020; Wilkinson et al., 2020) or spinal cord injury (López-Larraz et al., 2015). In addition to monitoring applications, these systems could also be integrated for home-based therapies, such as for motor rehabilitation (Bundy et al., 2017; Escolano et al., 2020), cognitive enhancement (Escolano et al., 2014), or closed-loop neurostimulation during sleep (Lee et al., 2020; Esparza-Iaizzo et al., 2021).

The terms of research applications, the fields that could most clearly benefit from this type of systems are those that involve out-of-the-lab brain monitoring. Remarkably, inexpensive and easy to wear EEG devices would enable large-scale recordings in ecological conditions (Dikker et al., 2017, 2021). This might constitute a breakthrough in the way we study the human brain (Matusz et al., 2019). On the one hand, brain activity could be measured in contexts where it is currently not possible, such as during daily life activities. On the other hand, the resources currently required to execute a scientific study with few dozens of participants, might allow to records hundreds or thousands of them.

In addition to clinical and research applications, brain monitoring with garments could be easily adopted for neuroscience applications in education, wellness, sports or industrial environments, incorporating these measurements to the range of bio-signals that are currently acquired by smart watches, rings, or chest-bands (Peake et al., 2018; Cannard et al., 2020; Di Pasquale et al., 2022).

## Conclusions

This paper presented the first proof of concept of a garment to measure EEG activity. We compared the Garment-EEG with a state-of-the-art Ag/AgCl dry EEG system in terms of spontaneous and evoked EEG activity, artifacts, skin-electrode impedance, and comfort. Under favorable recording conditions (low level of artifacts), the Garment-EEG provided comparable measurements to the metal-based EEG system, although it is more prone to getting affected by artifacts in adverse conditions, due to poorer contact impedances.

One of the most relevant advantages of this innovation is that it opens the door to creating EEG technology that can be broadly adopted by the general population, due to its comfort and its reduced manufacturing cost. This could allow large scale recordings for clinical and non-clinical purposes in ecological conditions.

An important final remark about the opportunities that this technology could enable is that this paradigm shift in brain monitoring should not take place without paying sufficient attention to its inherent risks. Experts in neuroethics are already working on creating recommendations for researchers, manufacturers and regulators to facilitate a responsible development of the neurotech field and its safe integration in our daily lives (Yuste et al., 2017).

## Author contributions

EL-L and JM conceived and designed the study. EL-L, CE, AR, LM and AA prepared the technology and the experimental setup. EL-L and AR participated in data collection. EL-L and CE analyzed the data, with inputs from the rest of the authors. EL-L and JM drafted the manuscript, with inputs from the rest of the authors. All the authors revised and approved the manuscript.

## Acknowledgments

The authors thank Diego Bayona for his support with data collection.

## Data availability statement

The raw data recorded with the Garment-EEG and the Dry-EEG systems will be shared as supplementary materials.

